# Selective Ablation of Cancer Cells with Low Intensity Pulsed Ultrasound

**DOI:** 10.1101/779124

**Authors:** David R. Mittelstein, Jian Ye, Erika F. Schibber, Ankita Roychoudhury, Leyre Troyas Martinez, M. Houman Fekrazad, Michael Ortiz, Peter P. Lee, Mikhail G. Shapiro, Morteza Gharib

**Affiliations:** Division of Engineering and Applied Sciences, California Institute of Technology, Pasadena, CA; Department of Immuno-Oncology, Beckman Research Institute, City of Hope, Duarte, CA; Division of Chemistry and Chemical Engineering, California Institute of Technology, Pasadena, CA

**Author notes:** address correspondence to co-senior authors M.G.S. or M.G.

## Abstract

Ultrasound can be focused into deep tissues with millimeter precision to perform non-invasive ablative therapy for diseases such as cancer. In most cases, this ablation uses high intensity ultrasound to deposit non-selective thermal or mechanical energy at the ultrasound focus, damaging both healthy bystander tissue and cancer cells. Here we describe an alternative low intensity pulsed ultrasound approach that leverages the distinct mechanical properties of neoplastic cells to achieve inherent cancer selectivity. We show that when applied at a specific frequency and pulse duration, focused ultrasound selectively disrupts a panel of breast, colon, and leukemia cancer cell models in suspension without significantly damaging healthy immune or red blood cells. Mechanistic experiments reveal that the formation of acoustic standing waves and the emergence of cell-seeded cavitation lead to cytoskeletal disruption, expression of apoptotic markers, and cell death. The inherent selectivity of this low intensity pulsed ultrasound approach offers a potentially safer and thus more broadly applicable alternative to non-selective high intensity ultrasound ablation.

## INTRODUCTION

High intensity focused ultrasound (HIFU) is a non-invasive therapeutic modality used clinically for tumor ablation [1–5]. By producing local hyperthermia and destructive cavitation [6], HIFU induces cell lysis, increases chemotherapeutic uptake [4, 7], and stimulates systemic anti-tumor immune responses [5, 8]. However, high intensity (I_SPTA_ >100 W/cm^2^) and high pressure (>10 MPa) focused ultrasound indiscriminately destroys healthy tissue as well as tumors [9, 10]. Consequently, safely implementing HIFU often requires costly MRI targeting [11] and is challenging in cancers near or invading into critical tissue [3].

Several approaches aim to increase ultrasound’s specificity. Molecularly targeted contrast agents, such as microbubbles [12, 13] locally amplify ultrasound’s disruptive effects, but are challenging to deploy in tumors due to the agents’ poor extravasation [14]. An alternative approach involves low intensity pulsed ultrasound (LIPUS). Low intensity (I_SPTA_ <5 W/cm^2^) and low frequency (<1 MHz) pulsed ultrasound produces mechanical effects without hyperthermia, resulting in neurostimulation [15], chemotherapy uptake [16], and bone repair [16–18]. However, its ability to selectively ablate cancer cells has not been studied, and its mechanisms of action are not fully understood [19–21].

Here we test the hypothesis that biomechanical differences between cancerous and healthy cell types cause these cells to have different responses to LIPUS, allowing selective ablation of cancer cells with targeted ultrasound waveforms. This hypothesis is predicated on cancer cells’ altered cellular/nuclear morphology, DNA content, nuclear-nucleolar volume ratios, cytoskeletal stiffness, and viscoelastic properties [22–25]. In computational studies, these differences were predicted to result in the differential response of malignant cells to specific ultrasound parameters when compared to healthy cells [26]. We use a cell suspension model to test this experimentally and examine the underlying acoustic and biophysical mechanisms.

## RESULTS

### Tuning frequency and pulsing of LIPUS allows for cancer selective cytodisruption

To test the hypothesis that LIPUS can selectively ablate cancer cells, we applied LIPUS to suspensions of human K562 and U937 cancer lines and primary T cells isolated from human peripheral blood mononuclear cells (PBMC), chosen as representative malignant and healthy cell types. Cell suspensions were placed in acoustically transparent-bottomed 24-well plates and insonated with a focused ultrasound transducer positioned in a water bath below (Fig 1a). We used pulsed ultrasound (10% duty cycle) with peak negative pressure (PNP) <1.2 MPa and I_SPTA_ <5 W/cm^2^. We confirmed that 60 seconds LIPUS at 0.67 MHz, 20ms pulse duration (PD) induced significant and irreversible cytodisruption of K562 cells at PNP > 0.6 MPa, as measured with ethidium homodimer-1 (Ethd-1) uptake (Fig 1b). Heating was always <1°C (**Fig Sup 1**). For further experiments, we selected 0.7 MPa PNP, which induced moderate cytodisruption.

**Figure 1.**
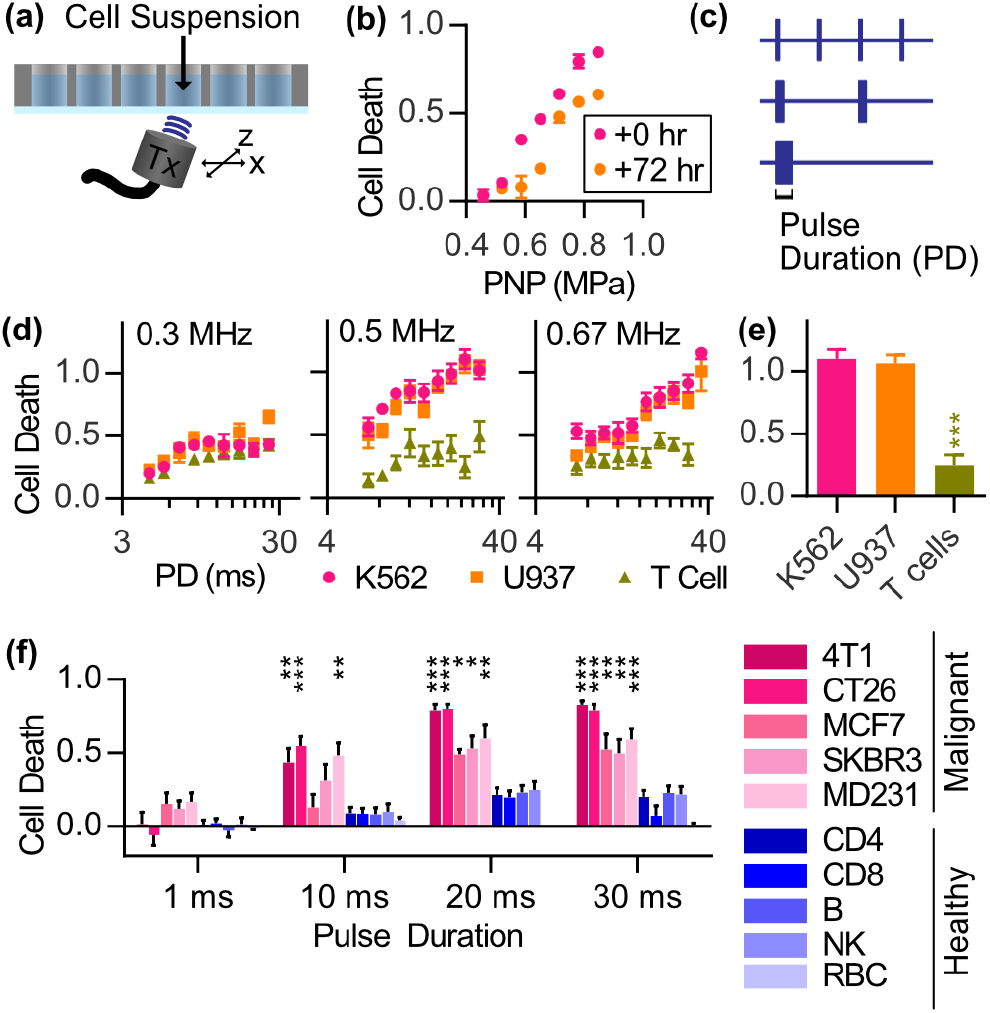
Screening reveals ultrasound parameters that induce cancer-cell selective cytodisruption. **a**, Schematic of mylar-bottom 24-well plate over water bath containing focused ultrasound transducer. **b**, K562 cell death (N=3, error bars SEM) in response to 0.67 MHz, 20ms PD, 10% duty cycle, 60 seconds, LIPUS at various peak negative pressures (PNP). 0.7 MPa PNP selected for future experiments. **c**, Diagram depicting constant energy while sweeping PD. **d, 0.**3 (N=4), 0.5 (N=9), 0.67 (N=9) MHz LIPUS induces frequency-, PD-, and cell-dependent cytodisruption. **e**, 0.5 MHz, 20ms PD LIPUS induces significantly less cell death (N=9, p<0.001) on T cells compared to either K562 or U937. **f**, 0.5 MHz LIPUS induced cancer-selective cell death (N=9) in mixed sample of healthy PBMC and cancer models (Table 1), measured through cytometry. RBC death assessed using hemoglobin release. Significance indicated as largest p-value from 2-tailed t-test between each cancer and each healthy cell model. (* p<0.05, ** p<0.01, *** p<0.001).

To test different pulsing patterns, PD and pulse repetition frequency were varied simultaneously to maintain constant energy (Fig 1c). We swept PD from 2-40ms with each of the 0.3, 0.5, and 0.67 MHz transducers (Fig 1d). We observed that cytodisruption was highly dependent on frequency, PD, and cell type. 20ms PD and 0.5 MHz maximized the selectivity with near complete cytodisruption for K562 and U937 and >80% survival for T cells (Fig. 1e). We found that cytodisruption increases with longer PD, despite the same total energy applied. Next, we assessed a broader panel of cell types at 0.5 MHz, using cancer cells in co-culture with PBMC (Table 1). The cancer cell models showed significantly more cytodisruption than subpopulations within PBMC at >10ms PD. (Fig 1f). RBCs exhibited virtually no disruption, as measured by hemoglobin leakage, under any condition.

**Table 1.** Cell Models Tested at Targeted Parameters

|  |  |  |
| --- | --- | --- |
| 4T1 | Mouse Cell Line | Mammary Gland, Epithelial (human breast cancer) |
| CT26 |  | Colon, Fibroblast (Carcinoma) |
| MCF7 | Human Cell Line | Mammary Gland, Epithelial (adenocarcinoma) |
| SK-BR-3 |  |  |
| MDA-MB-231 |  |  |
| CD4 | Human Primary Cells, Peripheral Blood Cells | CD3+, CD4+ |
| CD8 |  | CD3+, CD8+ |
| B |  | CD3-, CD19+ |
| NK |  | CD3-, CD19-, CD56+ |
| RBC | Bovine Primary Cells | Peripheral Blood Cells |

### LIPUS cytodisruption associated with cytoskeletal damage and activation of apoptotic, immunogenic cell death pathways

To characterize LIPUS cytodisruption’s biomolecular mechanisms, we evaluated CT-26 cells 2 days after 2-minute treatment with 0.5 MHz, 0.7 MPa LIPUS with flow cytometry. At >10ms PD, increased cell death and apoptosis was observed [27] (Fig 2a). Also, cells expressing calreticulin, a pro-phagocytic signal [28], increased while proliferative marker Bcl-2 [29] decreased (Fig 2b). The activation of apoptotic and phagocytic pathways may enhance LIPUS’ effectiveness as an anti-cancer therapy by promoting anti-tumor immune response.

**Figure 2.**
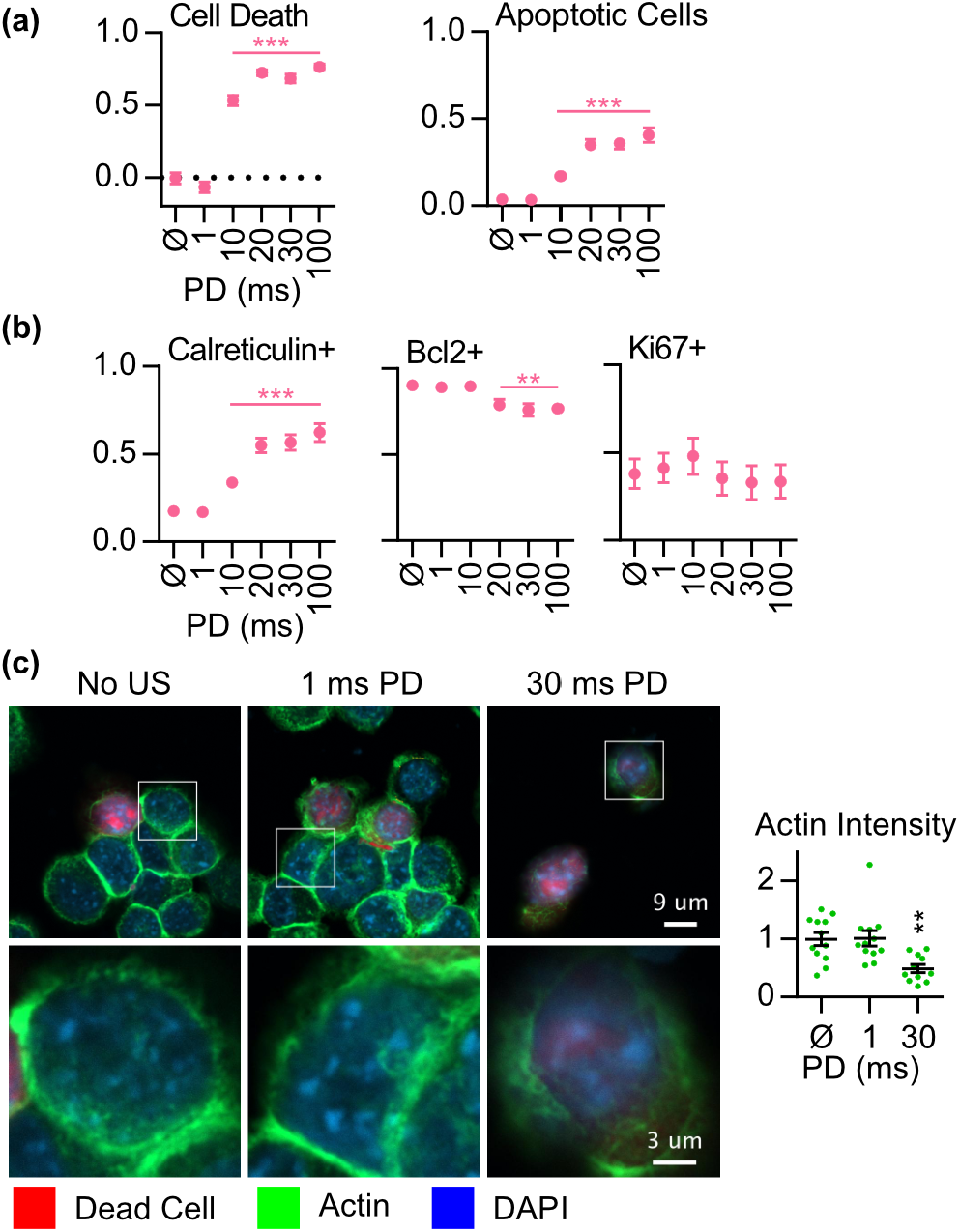
Ultrasound cytodisruption associated with apoptotic and pro-phagocytic pathways. **a**, CT-26 cells assessed 2 days after LIPUS (0.5 MHz, 0.7 MPa 20ms PD, 10% DC, 2 min). Fraction of surviving cells and apoptotic cells assessed using flow cytometry Ethd-1 vs Annexin V graphs (**Fig Sup 2)**. X-axis from no ultrasound (∅) to 1-100ms PD. Significantly increased cell death and apoptosis with >10ms PD LIPUS (N=12, error bars SEM). **b**, Increase in pro-phagocytic marker calreticulin (N=12) and decreased survival marker Bcl2 (N=8) but no change in proliferation marker Ki67 (N=8) with >20ms PD US. **b**, Confocal microscopy of CT-26 cells immediately after LIPUS. 30 ms PD LIPUS disrupted actin ring and significantly decreased actin stain intensity (N=12). (** p<0.01, *** p<0.001).

To evaluate LIPUS’ effect on the cytoskeleton, we performed confocal microscopy on CT-26 immediately after LIPUS. The actin cytoskeleton, stained with phalloidin, is qualitatively and quantitatively disrupted after insonation with 30ms PD LIPUS. This agrees with literature that states that LIPUS disrupts the cellular cytoskeleton [30, 31]. We demonstrate the novel property that at 1ms PD LIPUS, the cytoskeleton appears unchanged from the negative control (Fig 2c).

### Standing waves are necessary for LIPUS cytodisruption

Next, we investigated the physical mechanisms transducing LIPUS into cellular effects. Literature suggests that acoustic standing waves affect the mechanical forces experienced by cells [32, 33]. Such waves can result from interference of an incident ultrasound wave with its reflection, forming a spatially static pattern of pressure nodes and anti-nodes.

The pressure profile in acoustic 24-well plates revealed a standing wave pattern near the water-air interface. To examine whether this plays a role in cytodisruption, we constructed an acoustically transparent cuvette, where standing waves could be optionally introduced using a reflector (Fig 3a). We found that cells insonated in the absence of standing waves (0.67 MHz, 100ms PD) did not show significant cytodisruption, while cells treated in the presence of a reflector reproduced the cytodisruption observed in the 24-well plate (Fig 3b). Doubling the PNP in the reflector-free configuration to match the maximal pressure at standing wave anti-nodes did not induce significant cytodisruption. This suggests that standing waves are mechanistically required for LIPUS cytodisruption.

**Figure 3.**
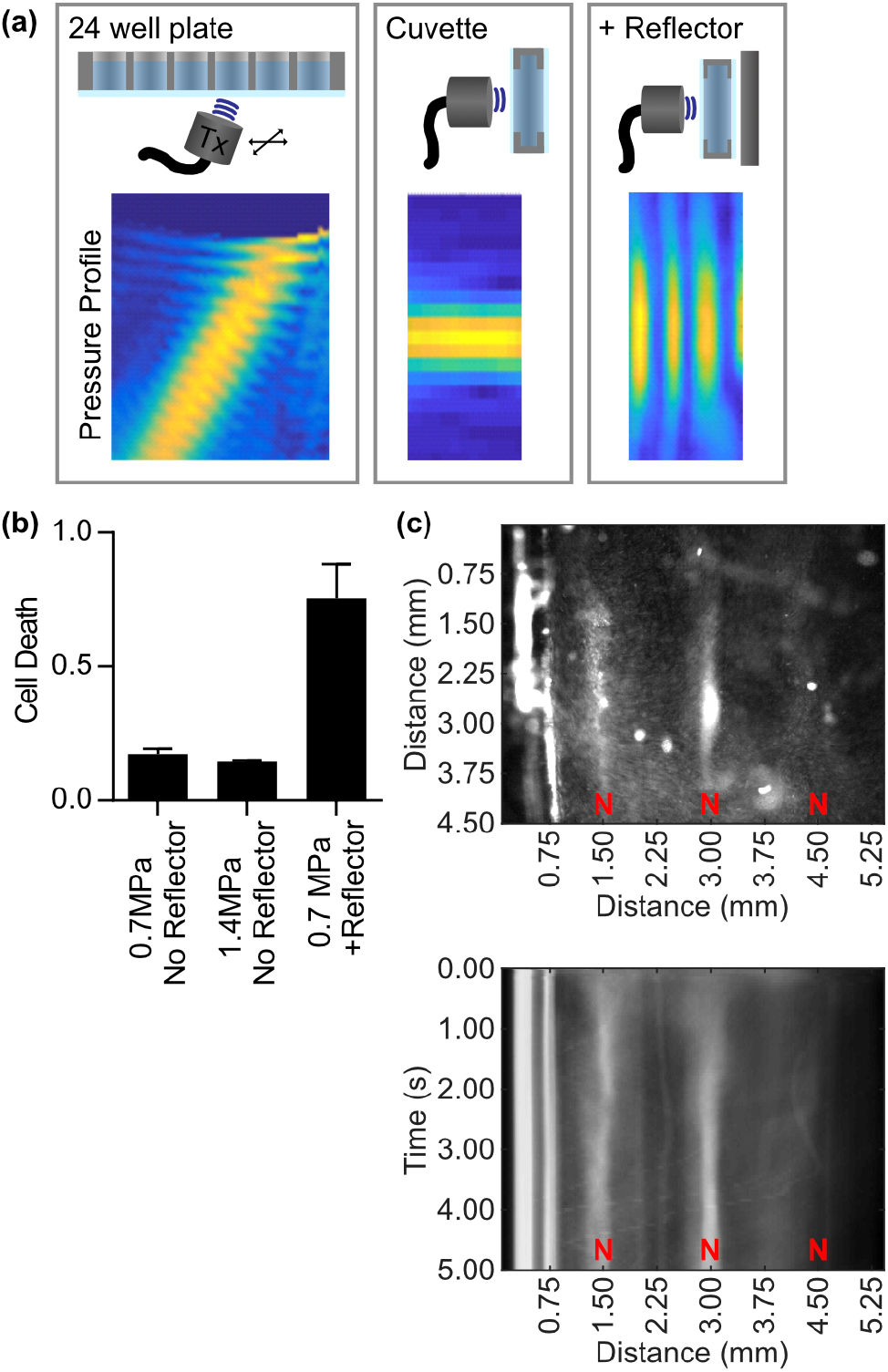
Standing waves required for LIPUS cytodisruptive effect. **a**, Schematics of experimental setups with pressure field measurements. Standing waves present in 24-well plate, but not in acoustic cuvette unless a metal reflector is introduced. **b**, 0.67 MHz, 100ms PD, 0.7 MPa LIPUS does not induce cell death in acoustic cuvette without reflector, even when doubling the pressure. However, when the reflector is added, cell death occurs as observed previously. **c**, Fluorescence microscopy demonstrates that cells accelerate toward nodes (N) in response to continuous 0.5 MHz, 0.7 MPa US. Upper image is still frame after 5 seconds of ultrasound, lower image represents average of y dimension (perpendicular to US) versus time. Cells achieve aggregation within 1 second.

Among other effects, acoustic pressure spatial gradients in standing waves give rise to acoustic radiation force that pushes cells toward pressure nodes [34]. We tracked the motion of fluorescently labeled K562 cells in response to continuous LIPUS in an imaging chamber and observed that 0.5 MHz ultrasound in a standing wave configuration propelled cells toward the nodes (Fig. 3c). A 100ms PD was not sufficient for cells to aggregate at nodes, which happened after ∼1 second under continuous ultrasound.

### Cell-mediated cavitation is required for LIPUS cytodisruption

Cavitation is a known mechanism for local amplification of acoustic pressure and cell killing [35]. To examine its role in LIPUS cytodisruption, we measured the acoustic emissions of cells treated with LIPUS in the acoustic cuvette using an orthogonally co-focused single-element passive cavitation detector (Fig 4a). We measured the signature of inertial cavitation (emissions with broad spectral content) and stable cavitation (harmonics of the transmitted frequency). As controls, we confirmed that no cavitation was measurable in degassed PBS cell buffer, while stable and inertial cavitation were detected from commercial Definity microbubbles (Fig. 4b).

**Figure 4.**
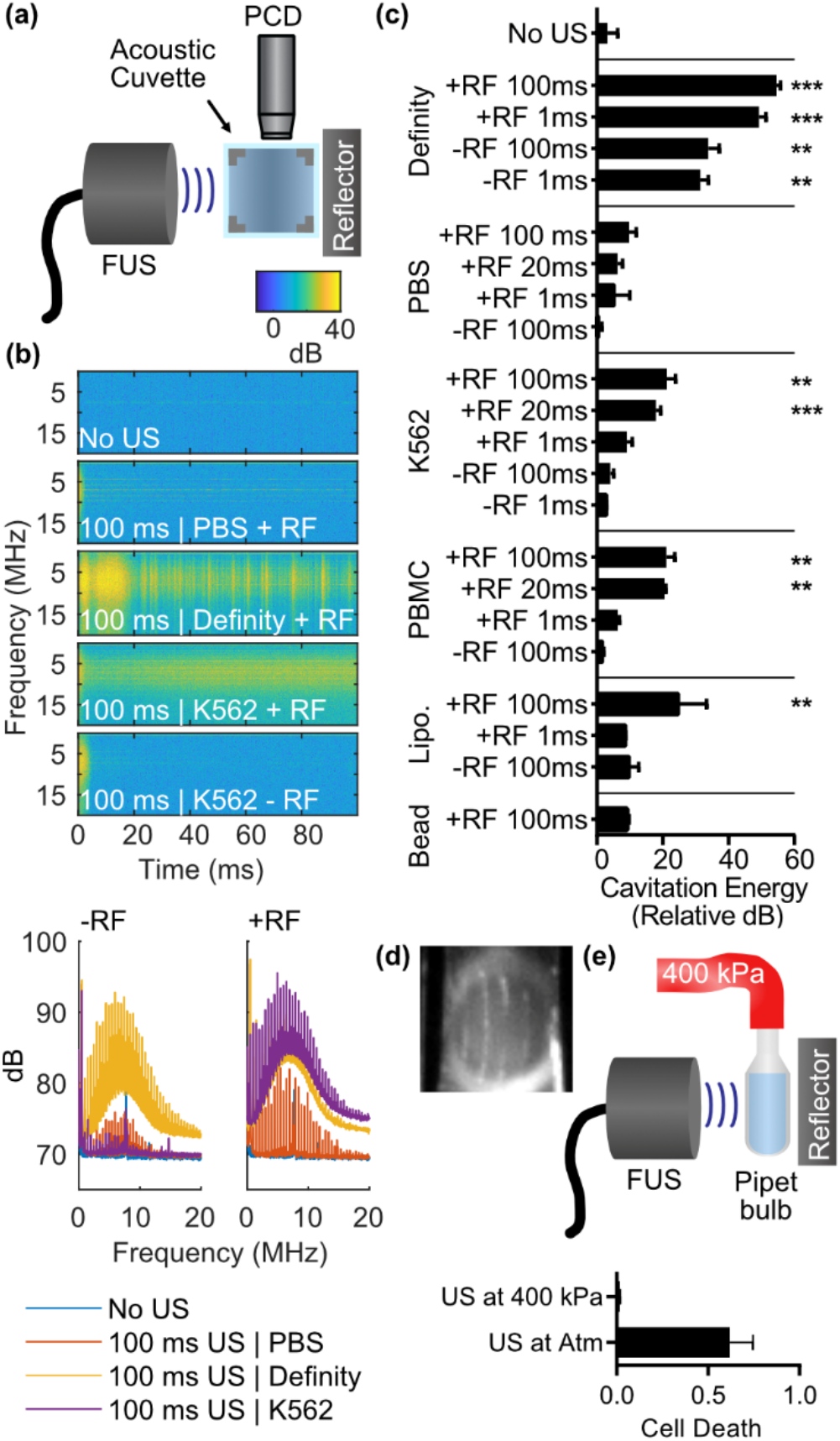
Cell-mediated cavitation is mechanistically necessary for cytodisruption. **a**, Schematic of passive cavitation detection setup using 10 MHz transducer orthogonally positioned to FUS transducer. **b)** Spectrogram of scattered signal from 100ms pulse of 0.5 MHz ultrasound transducer. Broadband signal of cavitation demonstrated with Definity positive control. No cavitation in PBS, however cavitation present in K562 suspensions only with reflector. **c)** Cavitation energy (∫ P^2^ dt) partially correlates with cytodisruption. Significant cavitation (compared to “No US”, ** p<0.01, *** p<0.001) observed with Definity, with K562 / PBMC with reflector (RF) and >20ms PD, and with liposomes with RF and 100ms PD. Note: PBMC cells induce cavitation, though they are resistant to LIPUS cyto-disruption. **d**, 100ms PD LIPUS with reflector induces cavitation bubbles formed in standing wave pattern in K562 sample. **e**, Schematic of pipet bulb pressurized to 400 kPa to form pressure chamber. At over-pressure, which suppresses cavitation, LIPUS cytodisruption is suppressed.

K562 cells generated both stable and inertial cavitation when exposed to 0.5 MHz, 100ms PD LIPUS in the presence of a reflector. The cavitation magnitude increased during the LIPUS pulse with an inflection point around 20ms (Fig 4b), a timescale similar to the PD needed for cytodisruption. No cavitation was seen without a reflector. Laser illumination of the cuvette revealed bubble formation in a standing wave pattern in response to long-PD ultrasound in the presence of the reflector (Fig 4d).

The conditions resulting in LIPUS-induced cavitation in K562 cell suspensions correlate with those causing cytodisruption, with both requiring standing-waves and PD >20ms (Fig. 4c). However, cavitation is not sufficient for cytodisruption, since PBMC, which are not strongly disrupted by tested LIPUS conditions, nevertheless produced similar amounts of cavitation (Fig 4c).

To determine which components of cell suspensions could be responsible for cavitation, we measured acoustic signal from LIPUS-exposed solutions of cell-sized 10 μm polystyrene beads and liposomes. No cavitation was detected from the beads. However, degassed liposomes did produce cavitation, suggesting that cell’s lipid contents may have a role in promoting cavitation (Fig 4c).

Finally, to confirm that cavitation is necessary for cytodisruption, we suppressed cavitation using over-pressure in an acoustically transparent pressure-chamber [36] comprising a plastic pipette bulb [37, 38] (Fig 4e). With the bulb pressurized to 400 kPa above ambient pressure, there was an almost complete reduction of cytodisruption of K562 cells in the chamber by 100ms PD LIPUS. This confirms that cavitation is mechanistically necessary.

### LIPUS results in translational motion of cells

To visualize LIPUS’ effect on target cells, we imaged K562 cells in suspension under transmitted laser illumination using an ultra-high speed camera. Cells were floating between two acoustically transparent films near an acoustic reflector generating standing waves. We recorded video at 5 Mfps starting 100ms after the beginning of insonation. We observed cells translating several microns along the axis of insonation at the ultrasound frequency, but not undergoing large-scale deformation **(Fig Sup 3, Sup Video)**. This suggests that either that LIPUS’ effect on cell shape are on the nanoscale and thus not detectable in this imaging paradigm or involve deformation or displacement of sub-cellular organelles relative to the cytoplasm.

### LIPUS cytodisruption attenuated in solid media

To investigate whether LIPUS cytodisruption occurs in cells embedded in solid media, we suspended K562 cells in agarose and acrylamide gels. We placed these gels in our acoustic cuvette with the reflector to generate standing waves **(Fig Sup 4a)**. Cell death was assessed using Ethd-1 fluorescence **(Fig Sup 4b)**. While statistically significant cytodisruption in agarose gels was observed, it was greatly attenuated compared to liquid suspensions **(Fig Sup 4c)**. This suggests that either the mechanical rigidity or the translational motion restriction imposed by a solid extra-cellular medium inhibits LIPUS cell killing.

## DISCUSSION

Our results demonstrate that specific parameters of LIPUS can induce cancer cell-selective cytodisruption. In an *in vitro* model, LIPUS applied at 0.5 MHz with a 20ms PD had the largest therapeutic margin in disrupting a diverse panel of cancer cells while leaving healthy blood and immune cells largely intact. PD >10ms, the formation of standing waves, and the emergence of cavitation were necessary to disrupt cancer cells. However, the presence of cavitation, which was seeded by cells and enhanced by standing wave ARF, was not sufficient to guarantee the disruption of any particular cell type. This suggests that while cavitation may locally amplify the pressure supplied LIPUS, a given cell type’s response to the resulting mechanical stress depends on its biophysical properties. This is consistent with the “oncotripsy” theory developed by Ortiz et al [26, 39], which suggests that cells respond to ultrasound at different resonant frequencies and with different fatigue behavior [40, 41].

Cancer-selective cytodisruption by LIPUS could fulfill the clinical need for safe non-invasive tumor ablation, complementing positional or molecular targeting approaches. Based on our results in suspension cell models, LIPUS applied within blood vessels could target blood cancer or circulating tumor cells [42]. The needed standing waves could be generated by engineering the acoustic field of one or more array sources or leveraging endogenous reflective surfaces such as bones. In addition, our data indicating that LIPUS-disrupted cells show markers of apoptotic and immunogenic cell death (ICD) suggest that disruption of cancer cells in circulation could stimulate immune responses against solid tumors elsewhere and strengthen the effect of conventional chemotherapeutic regimens [43–51]. Additional work is needed to extend this technology into the solid tumor context. While in our experiments cells in hydrogel phantoms responded weakly to LIPUS, this biochemical and mechanical context may not accurately represent the solid tumor milieu. In addition, even partial killing of solid tumor cells could be effective if it can precipitate an abscopal immune effect. Future *in vivo* studies are needed to test these hypotheses.

## MATERIALS AND METHODS

### High Throughput Ultrasound Experiments

Acoustically transparent 2.5 μm mylar film (Chemplex #100) was placed on the bottom of 24-well no-bottom plates (Greiner Bio-One #662000-06) that have been painted with a thin film of Sylgard 184 PDMS (Fisher Scientific # NC9285739) and heat treated at 60 °C for 24 hours. The resulting 24 well plates are watertight and have acoustically transparent bottoms. The plates were sterilized and loaded with cell samples as required for ultrasound exposure.

24 well plates were placed on a metal stage such that the mylar film was in contact with a water bath. One of the three available FUS transducers (0.3 MHz: Benthowave BII-7651/300, 0.5 MHz: Benthowave BII-7651/500, and 0.67 MHz: Precision Acoustics PA717) was attached to a metal arm angled 20 degrees from the normal of the water bath. A Velmex X-Slide motorized positioning system allowed the 3d motion of the arm allowing the transducers to be targeted at each well individually. The transducers were aligned using a Precision Acoustics fiber optic hydrophone to target the bottom center of well A1 on the 24 well plate. A MATLAB script controlled a signal generator (B&K #4054B) which generated a unique RF signal for each well of the plate and the Velmex positioning system. This signal was then amplified (AR #100A250B) and sent to drive the FUS transducers. The water bath was filled with distilled water which was degassed by a water conditioning system (ONDA #AQUAS-10) and heated to 37 degrees Celsius prior to experiments.

Fiber optic thermometry was used to measure the effect of insonation at the highest frequency 0.67 MHz and highest pressure 1.2 MPa PNP tested to confirm that no heating would occur in LIPUS experiments. Fiber was placed at ultrasound focus within acoustically transparent 24 well plate and temperature measurements were made for 1 ms and 100 ms pulse duration insonations. (**Fig. Sup 1**)

For the parameter search experiments (Fig. 1d) K562, U-937, or T cells were spun down and carefully resuspended in vacuum degassed PBS containing 2 μM ethidium homodimer-1 (Ethd-1) at 2 million cells in 2 mL PBS in each well of an acoustic 24 well plate. On each plate, 2 wells were loaded with 0.1% Triton X-100 as a positive control (pos) and 2 wells were un-insonated as a negative control (neg). Immediately after insonation, cell death for each well was estimated as Ethd-1 signal (s) as measured through plate reader as: cell death = (s_well_ − s_neg_) / (s_pos_ − s_neg_).

For the broad cell panel experiments (Fig. 1e), 2×10^6^ 4T1, CT26, MCF7, SK-BR-3 or MDA-MB-231 cancer cells were mixed with 2×10^6^ PBMCs in 2 mL degassed PBS respecitively and loaded into each well of an acoustic 24 well plate. After insonation, 2×10^4^ cells are cultured on 96 well plates for 2 days, and resuspended in PBS with 2 μM Ethd-1 prior to analysis with flow cytometry. For immune cell surface marker analysis, single-cell suspensions were stained with antibodies in PBS containing 2% fetal bovine serum. Antibodies to CD3(UCHT1), CD4(SK3), CD8(RPA-T8), CD19(SJ25-C1), CD33(P67.6) and CD56(5.1H11) were used to gate the CD4 T cells, CD8 T cells, B cells, myeloid cells and NK cells respecitively. Myeloid cells, which are largely undifferentiated cells with similar mechanical properties as cancer cells and comprise <1% of the PBMC cells, were excluded from analysis. Cell death for each subpopulation was determined from the count of cells that did not uptake Ethd-1 in comparison to untreated control.

Heparinized bovine red blood cells (Sierra for Medical Science), were diluted to 10% hematocrit in degassed PBS, then insonated as described above. After ultrasound, RBCs were centrifuged so that samples of supernatant could be assessed for hemoglobin release (Abcam ab234046). RBC death in resposne to LIPUS calculated as hemoglobin release compared to positive control 0.1% Triton X-100 and negative control of no ultrasound. (Fig. 1e)

### Biomolecular mechanism experiments

For CT26 cell apoptosis and proliferation marker analysis, 2 days after ultrasound treatment, CT26 cells were stained with anti-Calreticulin (Abcam) 30 min at room temperture. Annexin V binding buffer (Biolegend) was used for Annexin V staining. Fixation and permeabiliztion was performed with BD Cytofix/Cytoperm buffers (BD Biosciences) for Ki-67 and Bcl-2 intracellular antibodies staining. (Fig. 2a) Cell death was determined from the count of cells that did not uptake Ethd-1 in comparison to untreated control. The percentage of apoptotic cells reported is the fraction of Annexin V positive and Ethd-1 negative cells measured through flow cytometry. Cell signaling pathway markers Bcl-2 and Ki-67 were the fraction of Bcl-2 or Ki-67 stain positive and Aqua (fixable dead cell stain) negative cells. The pro-phagocytic marker calreticulin percentage reported was the fraction of calreticulin stain positive in all cells, live or dead.

For confocal experiments, CT-26 cells were allowed to settle on PDL-coated 1.5 μm coverslips for 1 hour. Slides were treated with fixable LIVE/DEAD stain (ThermoFisher #L34971), fixed in 4% paraformaldehyde and stained with phalloidin (Cayman #20549). Coverslips are then mounted with mountant containing DAPI (ThermoFisher #P36971). Four color confocal images were acquired with a 100x oil immersion objective at the Caltech Beckman Imaging Facility. (Fig 2b) To quantify, the actin signal intensity was measured on 12 cells imaged in each ultrasound treatment condition.

### Standing Wave Experiments

Pressure measurements performed using the fiber optic hydrophone positioned using the Velmex X-Slide. Color scale pressure maps represent peak negative pressure at each position within well or acoustic cuvette. Acoustic cuvettes were 1 cm × 1 cm 3d printed chambers with walls made of mylar film fixed with super glue. The cuvette was mounted to the bottom of the water tank and surrounded by distilled, degassed water. The center of cuvettes were aligned with the FUS transducer focus using fiber optic hydrophone and Velmex positioning system. A 3”x3”x0.5” rectangular prism block of aluminum was used as an acoustic reflector and positioned directly opposite from the transducer next to the cuvette (Fig 3a). K562 were loaded into the acoustic cuvette at 1 M cells / mL containing 0.2 μM Ethd-1. A 3d printed imaging chamber submersed in a water bath positioned the 0.5 MHz transducer a fixed position from an acoustic reflector, such that fluorescent imaging of a compartment containing GFP-labeled K562 at the focal point of the transducer could be achieved. Imaging was obtained through a 4x air objective (Olympus) (Fig. 3c).

### Cavitation Experiments

Using the same setup for standing wave experiments, a 10 MHz single element transducer (Olympus #U8421024) was positioned orthogonally to the FUS transducer, also aligned using the fiber optic hydrophone (Fig. 4a). Samples were loaded into the acoustic cuvette. Vacuum degassed PBS used as negative control. 10 μL of freshly resuspended Definity microbubbles (Lantheus Medical Imaging, Inc.) in degassed PBS used as positive control. Aluminum reflector used as described above to introduce standing waves. K562 or PBMC cells loaded at 1 M cells / mL in degassed PBS. Liposomes were generated from 14.0-18.0 PC Avanti Polar lipid suspended in chloroform which was lyophilized to remove chloroform, rehydrated using degassed 300 mOsm sucrose solution, sonicated for 10 minutes, heated at 40°C for 10 minutes, then degassed. Liposomes solution resuspended in degassed PBS to approximate lipid concentration in cell samples. 10 μM polystyrene beads at 1 M beads / mL also measured. Cavitation energy assessed in relative units by integrating the square of the pressure signal over time (Fig 4c). Image of cavitation bubbles taken using camera facing the single element transducer, with plane of laser illumination generated from laser light source and 1D diverging lens positioned above the acoustic cuvette (Fig 4d). A pressure chamber was constructed by attaching a compressed air line with a gauge pressure of 400 kPa onto an acoustically transparent plastic pipet bulb. Acoustic transmission through the pipet bulb was confirmed using hydrophone measurements. 1 M/mL K-562 cells loaded in degassed PBS containing 2 μM Ethd-1 into the pipet bulb. Cavitation from in samples loaded into the pipet bulb in place of the acoustic cuvette could be measured as described above. Cell death assessed using Ethd-1 signal (Fig 4e).

### Gel Experiments

Agarose gels were prepared by mixing 2% agarose in vacuum-degassed PBS at 65 °C with 2 M/mL K562 cells in PBS, adding 2 μM Ethd-1, and poured into 1 cm × 1 cm × 2.5 cm molds. Acrylamide gels were prepared as in this reference [52] with a final concentration of 1 M/mL K562 and 2 μM Ethd-1. Gels were insonated in the acoustic cuvette as described above. Cell death was calculated as magnitude of Ethd-1 signal observed at LIPUS focus on gel reader in comparison to signal from gels injected with 0.1% Triton X-100 and gels not treated with LIPUS.

### High Speed Camera Experiments

We assembled a high-speed microscopy setup capable of directly visualizing the effect of ultrasound on K562 cells. Our setup used a 2 W 532-nm laser (CNI, MLL-F532-2W) controlled by an optical beam shutter (Thorlabs SH05, KSC101). Right angle prism mirrors directed the laser light through a water bath and into a sample chamber containing the imaged samples. K562 cells were loaded into a custom-made acrylic cartridge containing an inner pocket surrounded by mylar film. A 3d printed holder positioned the cartridge such that the inner pocket was at the focus of the 0.5 MHz transducer. The cells were freely floating between two acoustically transparent films near an acoustic reflector that generate standing waves. A 100x water immersion Plan Fluor objective (Olympus) was used to image the target cells. A series of prism mirrors and converging lenses with focal lengths of 200 mm and 50 mm delivered the image into a Shimadzu HPV-X2 camera, which acquired 256 images over 51.2 µs, at a sampling rate of 5 million frames per second. To account for acoustic propagation through water, the camera was externally triggered to begin acquisition 100 ms after the start of the ultrasound pulse. A single pulse of 100 ms at 0.5 Mhz and 0.7 MPa PNP was used to insonate the sample in these experiments.

## Supporting information

High Speed Camera Video of Cell Response to LIPUS

## ACKNOWLEDGMENTS

The authors thank Sangjin Yoo, Di Wu, Avinoam Bar-Zion, Dan Piraner, and Mohamad Abedi for helpful discussion. We thank Sangjin Yoo for guidance with ultrasound LIPUS bioeffects in neuromodulation, Dan Piraner for cell culture and fluorescent protein transfection, Hunter Davis for input on HFR imaging optics, and Di Wu for fluorescent cell tracking experiments. We thank Michael R. Bailey for his assistance in designing the pipet bulb pressure chambers. We thank Maayan Harel (www.maayanillustration.com) for the illustrations in this paper. Confocal microscopy experiments were performed at Caltech’s Beckman Imaging Facility. This project was supported by Amgen CBEA 2017, 2018 and the Caltech/ City of Hope collaborative grant 2019. AR and SLT were supported by Caltech Summer Undergraduate Research Fellowships.

## AUTHOR CONTRIBUTIONS

DRM, MO, MGS, MG conceived the study. DRM, JY, AR, SLT, MO, PL, MGS, MG designed, planned, and conducted the experiments in this paper. DRM and JY analyzed the data. All authors discussed the results. DRM, JY, and MGS wrote the manuscript with input from all the authors. All the authors have given their approval for the final version of the manuscript. MGS and MG supervised the research.

## Competing interests

The authors declare no competing financial interests.

