## Supplemental Figures, Materials and Methods for "Selective Ablation of Cancer Cells with Low Intensity Pulsed Ultrasound"

### Supplementary Figures

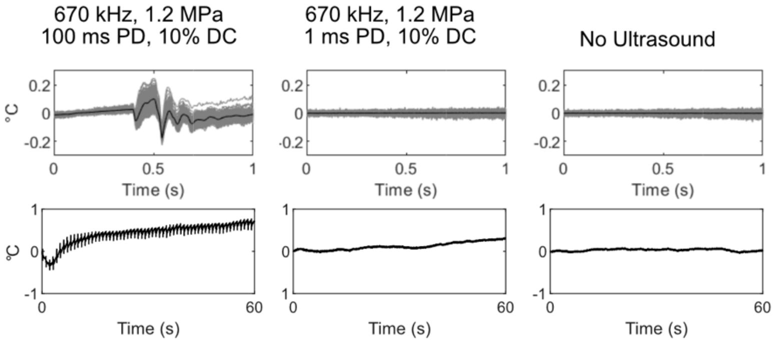


**Figure Sup 1 | No temperature change induced by LIPUS.** Assessed temperature change using fiber optic hydrophone system at focus of FUS transducer in 24-well plate configuration. LIPUS applied at 0.67 MHz, 1.2 MPa PNP, with 10% duty cycle did not induce any appreciable change (<1 °C) in temperature even with 100 ms pulse duration. Top row shows each temperature trace for each second in grey, with average change in black. Bottom row shows all data acquired in 60 seconds of LIPUS. Similar measurements for 300, 0.5 MHz transducers showed even less heating (not shown).

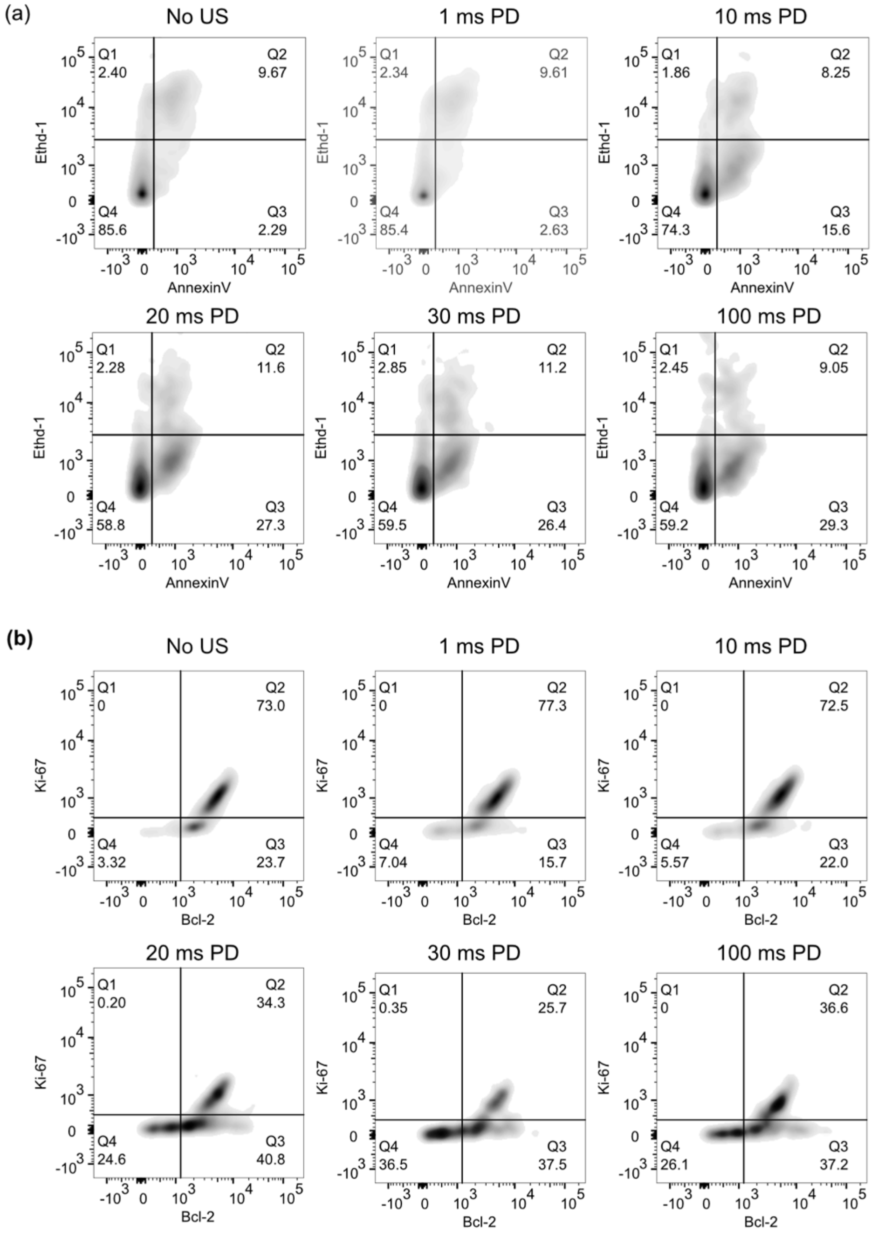


**Figure Sup 2 | Flow cytometry gates used for phagocytic, apoptotic analysis**. **a,** Ethidium homodimer (Ethd-1) vs AnnexinV stain generates the following quadrants. Q4: live cells, Q3: live cells undergoing apoptosis, Q2-Q1: dead cells. **b,** Ki-67 vs Bcl-2 stains used to identify Bcl2+ and Ki-67+ populations.

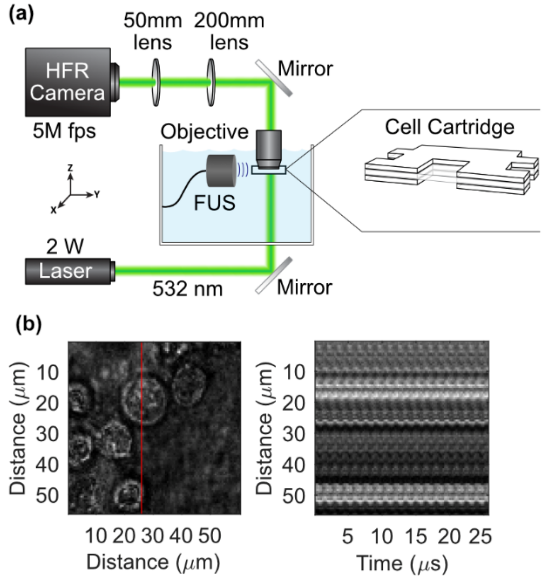


**Figure Sup 3** | **High speed video demonstrates no large-scale cell deformation during ultrasound insonation. a,** Schematic of high frame rate camera setup enabling cellular imaging at 5 Mfps. **b,** 1d trace over time demonstrates translation of cell of ~1 micron after 100ms of 0.67 MHz ultrasound exposure. Video in shows no visible deformation, only translation.

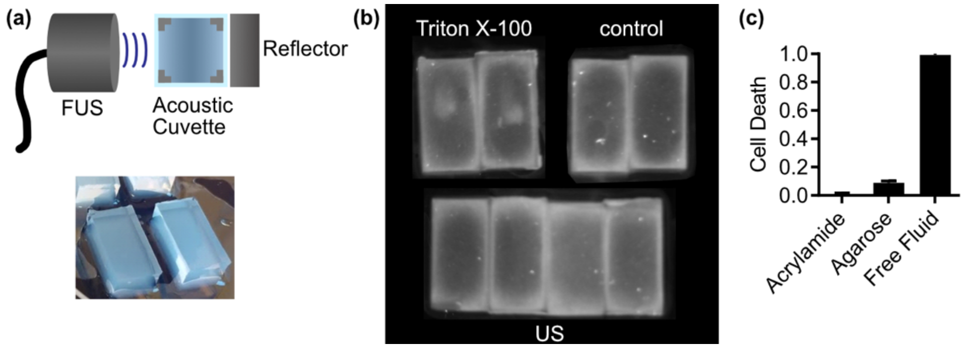


**Figure Sup 4** | **Cells suspended in gel medium undergo attenuated cytodisruption in response to our targeted LIPUS parameters. a,** Schematic diagram and picture demonstrating 1 cm thick 1% agarose gels containing K562 cells and their placement between the 0.5 MHz transducer and the metal reflector. **b,** Cytodisruption was assessed using ethidium homodimer assessed in gel reader, positive control had injection of Triton X-100 in center of gel. **c,** Cell death is completely attenuated in acrylamide gels compared to free fluid condition. Cell death is attenuated, but still significant in agarose gels. Observed using ethidium homodimer-1 fluorescence after ultrasound at 0.5 MHz, PD 100 ms.
